# Comprehensive variant and haplotype landscapes of 50,500 global SARS-CoV-2 isolates and accelerating accumulation of country-private variant profiles

**DOI:** 10.1101/2020.07.09.193722

**Authors:** Lishuang Shen, Jennifer Dien Bard, Jaclyn A. Biegel, Alexander R. Judkins, Xiaowu Gai

**Author notes:** Correspondence to: Xiaowu Gai, PhD, Center for Personalized Medicine, Division of Genomic Medicine, Department of Pathology and Laboratory Medicine, 4650 Sunset Blvd., Mailstop #173, Los Angeles, CA 90027.

## Abstract

Understanding the genetic etiology of COVID-19 requires a comprehensive understanding of the variant and haplotype landscape of all reported genomes of SARS-COV-2, the causative virus of the disease. Country-, state/region- and possibly even city-private variant profiles may contribute to varied disease exemplifications and fatality rates observed across the globe along with host factors such as age, ethnicity and comorbidity. The Children’s Hospital of Los Angeles (CHLA) COVID-19 Analysis Research Database (CARD) captures up-to-date fulllength SARS-CoV-2 sequences of ~50,500 isolates from GISAID, GenBank, CHLA Center for Personalized Medicine, and other sources (as of June 18, 2020). Among which, 49,637 isolates carry at least one variation from the reference genome NC_045512, a total of 6,070 variants and 2,513 haplotypes were detected in at least three isolates independently. Together, they constituted the most likely SARS-CoV-2 variant and haplotype landscapes world-wide currently.

Evidence supporting positive (orf3a, orf8, S genes) and purifying (M gene) selections were detected, which warrants further investigation. Most interestingly, we identified 1,583 country-private variants from 10,238 isolates (20.6% overall) reported in 48 countries. 807 country-private haplotypes, defined as a haplotype shared by at least 5 isolates all of which came from the same country, were identified in in 8,656 isolates from 39 countries. United Kingdom, USA, and Australia had 464, 166 and 32 private haplotypes respectively, comprising 22.4%, 16.6% and 16.4% of the isolates from each country. Together with their descendent and private haplotypes with fewer members, 22,171 (45.8%) isolates carried country-private haplotypes globally. The percentage were 28.2-29.6% in January to March, and rapidly increased to 46.4% and 59.6% in April and May, co-occurring with global travel restrictions. The localization of the variant profiles appeared to be similarly accelerating from 14.2% in March and 28.4% in April to over 40% isolates carrying the country-private variants around May.

In summary, a common pattern is seen world-wide in COVID-19 in which at the onset of disease there appeared to be a significant number of SARS-CoV-2 variants that accumulate quickly and then begin to rapidly coalesce into distinct haplotypes. This may be the result of localized outbreaks due to factors such as multiple points viral introduction, geographic separation and the introduction of policies such as travel restriction, social distancing and quarantine, resulting in the emergence of country-private haplotypes.

## Introduction

Comprehensive analysis of SARS-CoV-2 genomes is critically important to determine transmission patterns and identify haplotype and variant signatures that may be associated with evolving pathophysiology and virulence. The rapid accumulation and sharing of SARS-CoV-2 genome sequences at an unprecedented speed have greatly facilitated such efforts. Since the first SARS-CoV-2 genome sequence was reported in January of 2020, as of June 18th, there have been over 48,000 sequences deposited to GISAID (https://www.gisaid.org/; Elbe and Buckland-Merrett, 2017; Shu et al., 2017) and 8,000 sequences submitted to NCBI Virus (https://www.ncbi.nlm.nih.gov/genbank/sarscov-2-seqs/), the China National Center for Bioinformation (CNCB) 2019 nCoV Resource (https://bigd.big.ac.cn/ncov/; Zhao et al., 2020) and other data repositories. These data have been used to generate phylogenetic studies most of which have been country-private, while others have focused on global virus phylogenetic and transmission analyses. However, the full-scale characterization of SARS-CoV-2 genomes reported to date, at the time of this writing exceeding 50,000 worldwide, has not yet been reported.

In our previous study, we established a comprehensive COVID-19 genomic resource, CHLA CARD (https://covid19.cpmbiodev.net/), by harmonizing data from GISAID, NCBI Virus, CNCB and other resources (Shen et al., 2020). Using CHLA CARD, we performed a comprehensive analysis of all publically available SARS-CoV-2 genome sequences at the time of study. We called variants from each genome sequence, and annotated each isolate and variant with associated demographic data and functional predictions. Our categorical analyses of variants and haplotypes provided a snapshot of SARS-COV-2 viral genetics globally, six months into the COVID-19 pandemic. The functional and pathophysiological importance of these variants and haplotypes remains to be determined. However, based on the differential selections of variants in different genes, there is the potential for emergence of a number of different strains, at least some of which might differ in their transmissibility and virulence. Continuous monitoring of the country-private variant and haplotype profiles will lead to in-depth understanding of how the COVID-19 pandemic evolves from the viral genetic perspective, and will be critical as countries across the world move to re-open, often with variable degrees of control of COVID-19. Such an approach may hopefully aid in the timely identification of variants and haplotypes that are potentially associated with significant changes in transmissibility and virulence as they emerge.

## Results

### Overview of over 50K SARS-CoV-2 isolates

The merged SARS-CoV-2 collection as of June 18, 2020 consisted of ~50,500 isolates after removing duplicates among data sources. These were complete or near complete genome sequences (> 29,600 bases) with over 97% of the isolates originating from Asia, Europe and USA. Non-human host sequences were excluded. SARS-CoV-2 genome sequences of ~700 isolates were identical to the reference genome NC_045512.2. The remaining isolates carried at least one variant when compared against the reference, and were included in the subsequent variant and haplotype analyses. The number of variants per isolate ranged from 0 to over 30, and isolates with 30 or more variants were excluded from the analysis because they were regarded as likely low-quality sequences. The remaining 49,637 isolates belonged to 93 geographic locations; 22,068 (44.5%) isolates in the United Kingdom (UK) and 10,397 (20.9%) isolates in the US. Notably underrepresented were isolates from African (578) and South American (647) countries (**Table 1**). A total of 459 isolates were classified as imported based on the reported country of exposure being different to the country of the resident. Iceland had a highest number of imported isolates (198) followed by India (34) and China (30). A total of 17,100 variants and 24,633 potential haplotypes were detected from any isolate. However, the submitted genome sequences were generated by a variety of laboratories and sequencing technologies, not all of which provided the same or uniform coverage across the viral genome. To account for this variation, we further filtered the variants and haplotypes by requiring their presence in at least 3 isolates to be included in downstream analysis (**Table 1, 4**).

**Table 1.** Summary by geographic regions of variants and haplotypes present in at least 3 isolates.

| **Region** | **Variants** | **Region Isolates** | **Haplotypes** | **Region Isolates** | **Haplotype size** |
| --- | --- | --- | --- | --- | --- |
| Africa | 379 | 575 | 52 | 205 | 7 |
| Asia | 1593 | 3498 | 222 | 1275 | 7.5 |
| Europe- non-UK | 2219 | 9035 | 507 | 4445 | 8 |
| Europe- UK | 3649 | 22063 | 1313 | 11817 | 8 |
| North America - Non-USA | 458 | 921 | 88 | 484 | 6.5833 |
| North America- USA | 2481 | 10389 | 547 | 5025 | 8 |
| Oceania | 965 | 2430 | 215 | 1288 | 7.5714 |
| South America | 371 | 647 | 62 | 269 | 7 |

**Table 2.**
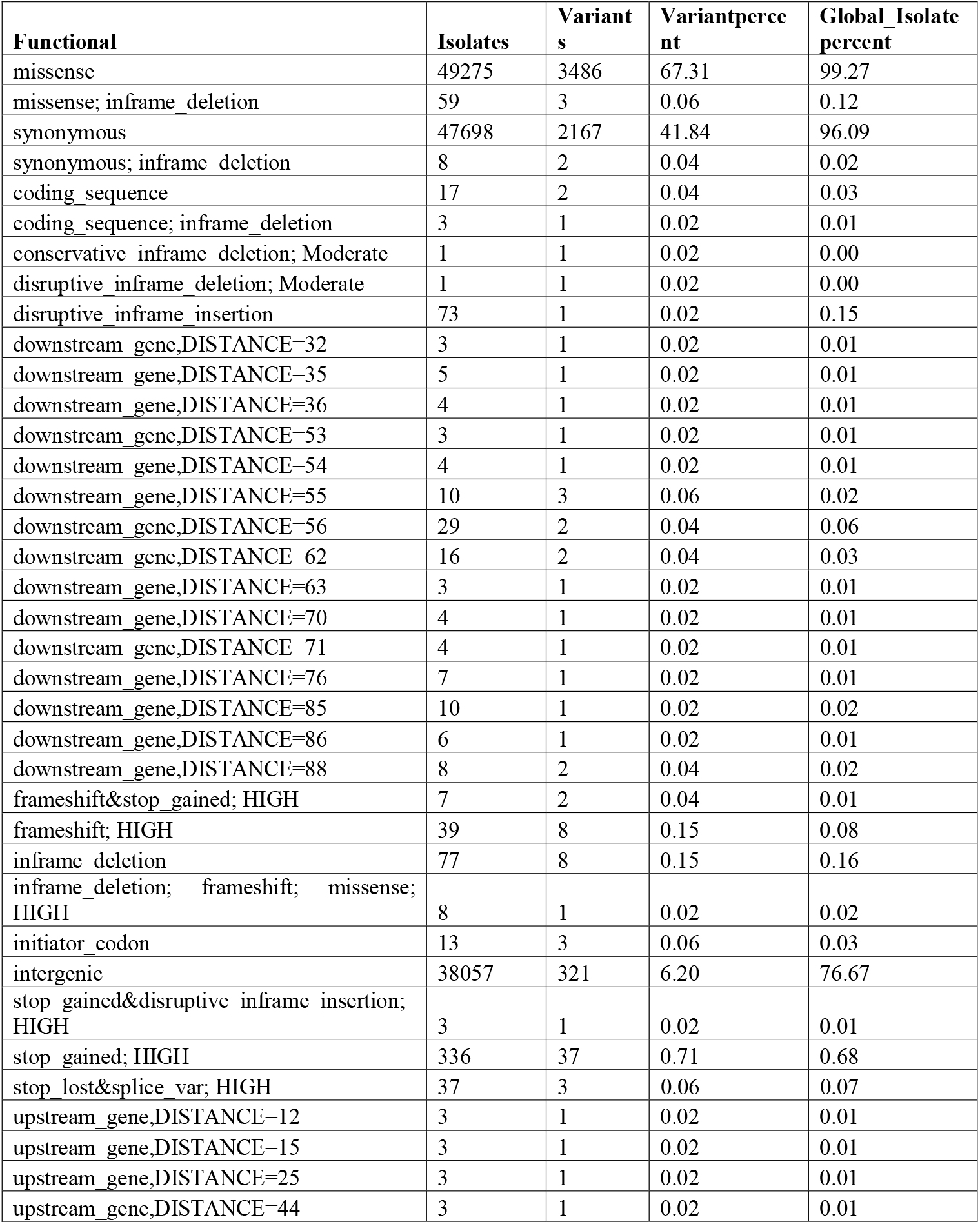
Functional prediction summary of variants present in 3 or more isolates.

| **Functional** | **Isolates** | **Variant**<br>**s** | **Variantperce**<br>**nt** | **Global_Isolate**<br>**percent** |
| --- | --- | --- | --- | --- |
| missense | 49275 | 3486 | 67.31 | 99.27 |
| missense; inframe_deletion | 59 | 3 | 0.06 | 0.12 |
| synonymous | 47698 | 2167 | 41.84 | 96.09 |
| synonymous; inframe_deletion | 8 | 2 | 0.04 | 0.02 |
| coding_sequence | 17 | 2 | 0.04 | 0.03 |
| coding_sequence; inframe_deletion | 3 | 1 | 0.02 | 0.01 |
| conservative_inframe_deletion; Moderate | 1 | 1 | 0.02 | 0.00 |
| disruptive_inframe_deletion; Moderate | 1 | 1 | 0.02 | 0.00 |
| disruptive_inframe_insertion | 73 | 1 | 0.02 | 0.15 |
| downstream_gene,DISTANCE=32 | 3 | 1 | 0.02 | 0.01 |
| downstream_gene,DISTANCE=35 | 5 | 1 | 0.02 | 0.01 |
| downstream_gene,DISTANCE=36 | 4 | 1 | 0.02 | 0.01 |
| downstream_gene,DISTANCE=53 | 3 | 1 | 0.02 | 0.01 |
| downstream_gene,DISTANCE=54 | 4 | 1 | 0.02 | 0.01 |
| downstream_gene,DISTANCE=55 | 10 | 3 | 0.06 | 0.02 |
| downstream_gene,DISTANCE=56 | 29 | 2 | 0.04 | 0.06 |
| downstream_gene,DISTANCE=62 | 16 | 2 | 0.04 | 0.03 |
| downstream_gene,DISTANCE=63 | 3 | 1 | 0.02 | 0.01 |
| downstream_gene,DISTANCE=70 | 4 | 1 | 0.02 | 0.01 |
| downstream_gene,DISTANCE=71 | 4 | 1 | 0.02 | 0.01 |
| downstream_gene,DISTANCE=76 | 7 | 1 | 0.02 | 0.01 |
| downstream_gene,DISTANCE=85 | 10 | 1 | 0.02 | 0.02 |
| downstream_gene,DISTANCE=86 | 6 | 1 | 0.02 | 0.01 |
| downstream_gene,DISTANCE=88 | 8 | 2 | 0.04 | 0.02 |
| frameshift&stop_gained; HIGH | 7 | 2 | 0.04 | 0.01 |
| frameshift; HIGH | 39 | 8 | 0.15 | 0.08 |
| inframe_deletion | 77 | 8 | 0.15 | 0.16 |
| inframe_deletion; frameshift; missense; HIGH | 8 | 1 | 0.02 | 0.02 |
| initiator_codon | 13 | 3 | 0.06 | 0.03 |
| intergenic | 38057 | 321 | 6.20 | 76.67 |
| stop_gained&disruptive_inframe_insertion; HIGH | 3 | 1 | 0.02 | 0.01 |
| stop_gained; HIGH | 336 | 37 | 0.71 | 0.68 |
| stop_lost&splice_var; HIGH | 37 | 3 | 0.06 | 0.07 |
| upstream_gene,DISTANCE=12 | 3 | 1 | 0.02 | 0.01 |
| upstream_gene,DISTANCE=15 | 3 | 1 | 0.02 | 0.01 |
| upstream_gene,DISTANCE=25 | 3 | 1 | 0.02 | 0.01 |
| upstream_gene,DISTANCE=44 | 3 | 1 | 0.02 | 0.01 |

**Table 3.**
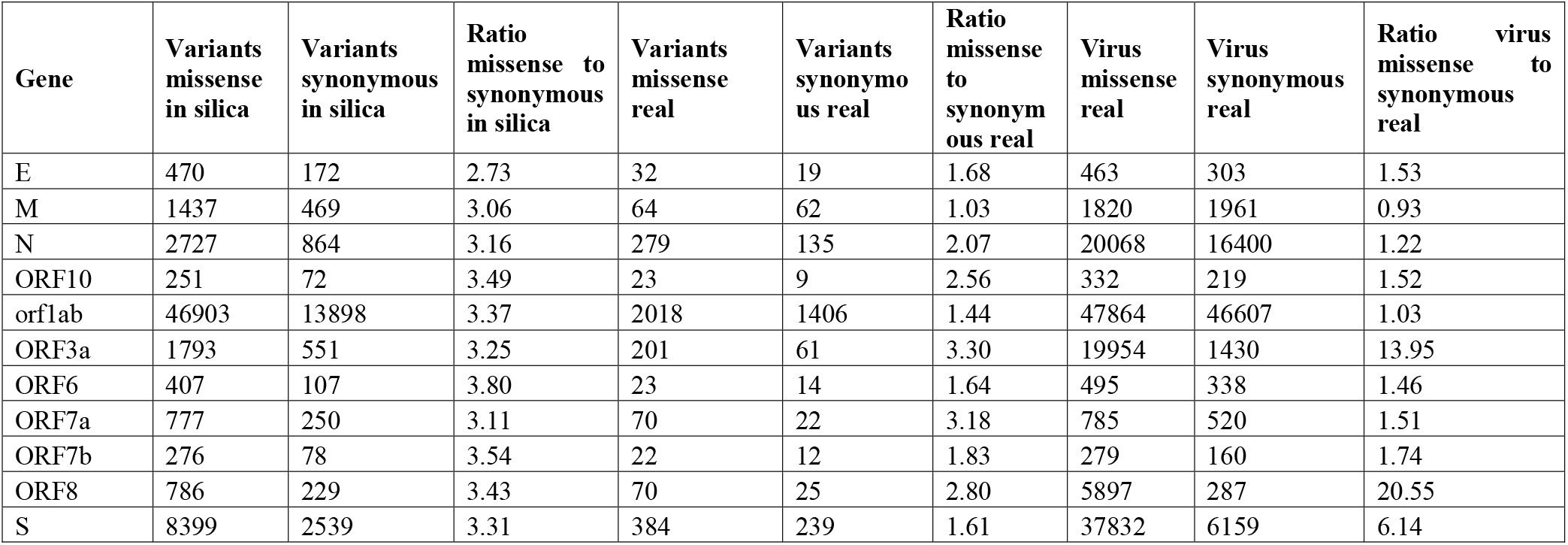
Per gene functional prediction summary of missense and synonymous variants present in 3 or more isolates.

**Table 4.**
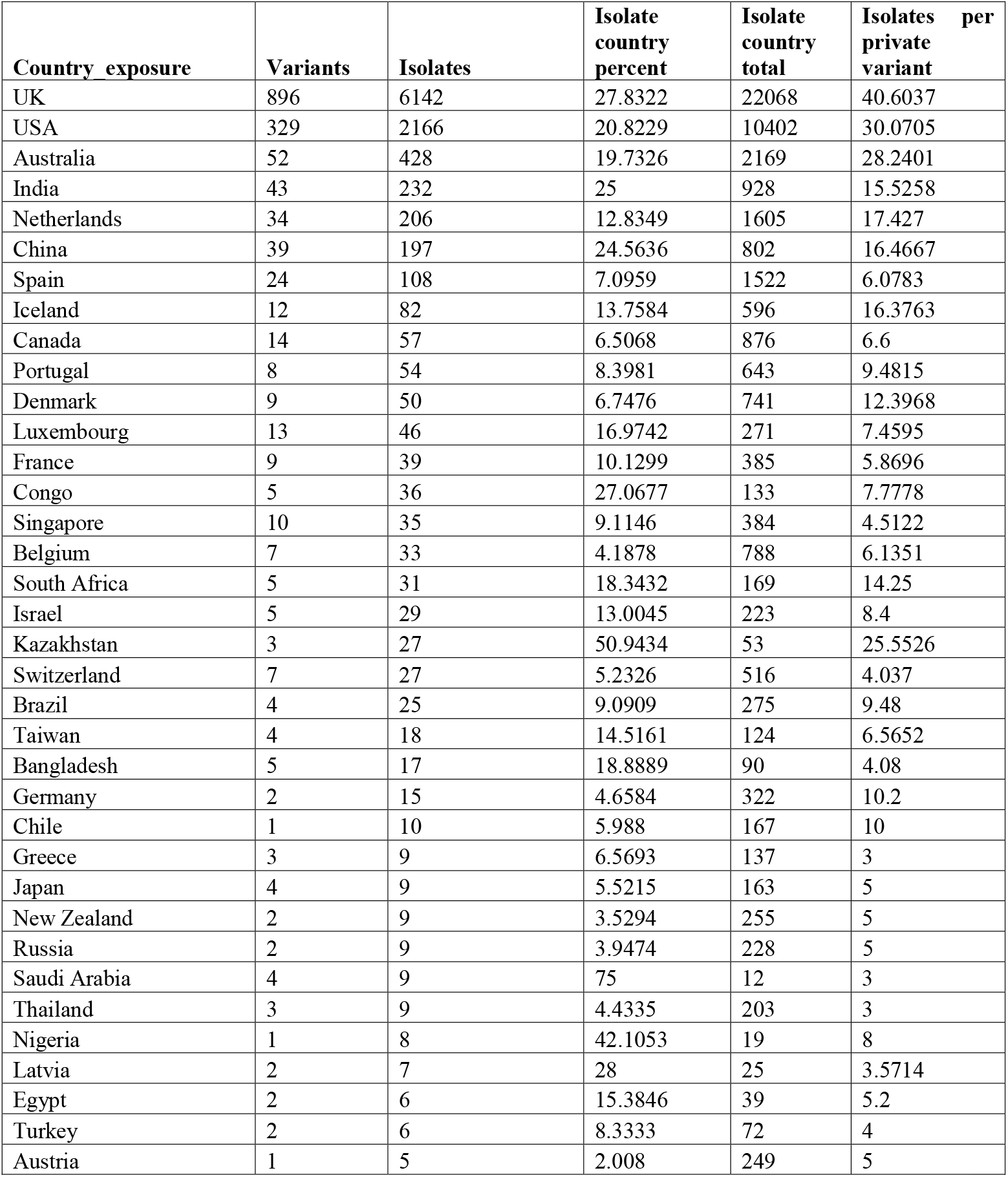
Country-private variants present in 3 or more isolates.

### High-quality variants and haplotypes

A total of 6,070 variants and 2,513 haplotypes were identified in at least three global isolates independently and were deemed reliable and less likely to be sequencing artifacts (**Table 1**). In total, these 6,070 variants included 3,486 missense, 2,167 synonymous, 336 intergenic, 12 inframe deletions/insertion, 10 frameshift, 41 stop gained/lost, and several other noncoding variants (**Table 2**). The number of missense variants was significantly higher than that of synonymous variants, overall across all genes, and for each individual gene. The number of missense variants was 77% more than synonymous variants but less than the *in silica* predicted ratio of ~3.0 based on random chance. The gene level numbers were 279 vs 135 (2.0x) in N gene, and 384 vs 239 (1.6x) in S gene for the global isolates (**Table 3**). The four most common mutations (241-C-T/5UTR-orf1ab, 3037-C-T/orf1ab:p.Phe924Phe, 14408-C-T/orf1ab:p.Pro4715Leu, 23403-A-G/S:p.Asp614Gly) were each present in 70.3%-71.8% of non-US isolates and 67.5%-69.3% of US isolates (**Figure 1**). The multi-nucleotide variant “28881-G-A, 28882-G-C, 28883-G-A” causing a 2-amino acid variant N:p.Arg203_204LysGly -> N:p.Arg203_204LysArg, was present in 32.0% of the non-US isolates, with a significant presence in isolates from the UK (41.3%), other European countries (24.0%) and Australia (13.5%), but not in the isolates from the USA (3.4%) (**Figure 1**).

**Figure 1.**
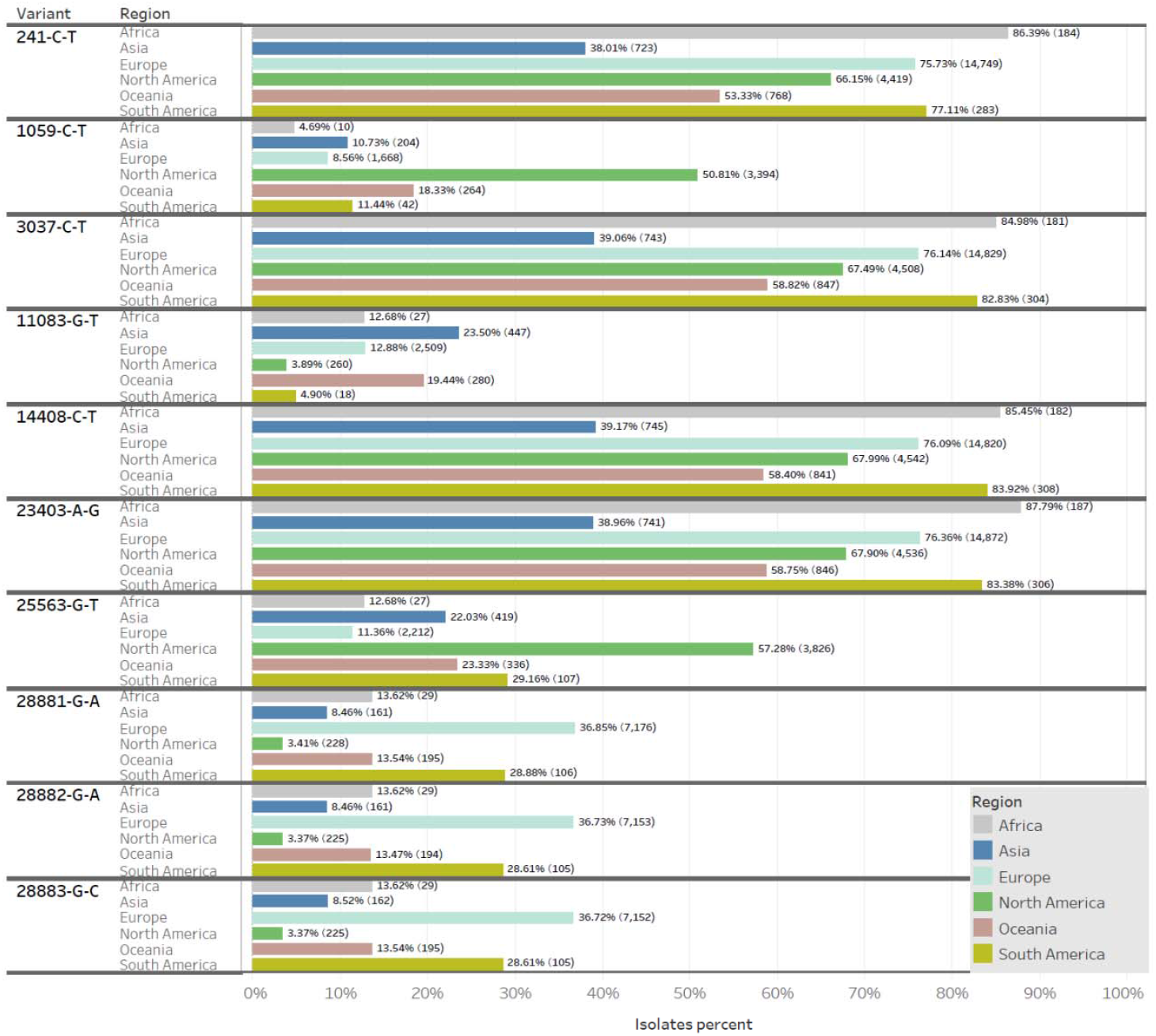
The continental-level percentages of isolates carrying the most abundant variants those were present in at least 3,000 isolates. **Legend: The number of isolates per continent are enclosed in parentheses.**

Across geographic locations, the total number of variants was highest in the UK isolates (3,649), followed by USA (2,481), while Asia (1,593) and Oceania (965) collections had far fewer variants. It is worth noting, however, that rates of sequencing vary significantly across these geographies.

The largest three haplotypes were the 1,771-isolate haplotype “241-C-T, 3037-C-T, 14408-C-T, 23403-A-G, 28881-G-A, 28882-G-A, 28883-G-C”, the 1,458-isolate haplotype “241-C-T, 1059-C-T, 3037-C-T, 14408-C-T, 23403-A-G, 25563-G-T”, and the 727-isolate haplotype “241-C-T, 3037-C-T, 14408-C-T, 23403-A-G”. They were all predominantly present in Europe and USA.

### Enrichment and depletion of missense variant-carrying isolates and selection

The missense variants predicted *in silica* based on random chance were about 3 times that of synonymous variants per gene, in the range of 2.7~3.8 (**Table 3**). The observed ratios of variant ranged from 1.03 for M gene, to 3.30 for ORF3a gene, but all the ratios were lower than the *in silica* predicted ratios based on random chance. The ratio of isolates with at least one missense variant in a gene versus isolates with at least one nonsynonymous variant in the same gene, was 0.93 in the M-gene (Chi-squared test p-value of depletion < 2.2e-16), suggesting the possibility that this gene was undergoing strong purifying selection. Purifying selection could have affected the M gene whose missense variant–carrying isolates were significantly less frequent than expected from the in-silica predicated variant types. In contrast, the ratios of isolates were 20.6 in ORF8, 14.0 in ORF3a gene, and 6.1 in S-gene with the Chi-square p-values of enrichment for missense variants all below 2.2e-16, suggesting the possibility that these genes may have been under positive selection for missense variants. Further analysis including functional validation would be required to confirm the significance of these changes. In-depth analyses that take into consideration of emergence timing, geographic location, founder effect, and ancestral haplotype are needed before any conclusion can be reached. Ultimately, questions about the fitness of variants require *in vitro* functional studies. There have been speculations that some super-contagious lineages of SARS-CoV-2 may be evolving towards higher transmissibility. 25563-G-T (ORF3a:p.57Q>H) is a noteworthy candidate. It emerged first in France on 2/21/2020, later in USA on 02/27/2020 (**Figure 3**), and quickly dominated the USA isolates since 03/02/2020, present in 60.0% of isolates. It also appeared in 22% and 11.4 % of isolates from Asia and European union, but was rare in the UK. In France, the frequency of this allele had continued to change and the allele was rare in isolates since mid-April. Tracing this allele in the context of haplotype evolution may help clarify whether it was imported or spontaneously occurred again in USA isolates. The 23403A>G (S:p:614D>G) mutation which quickly dominated USA and Europe caused 4-time increase of higher transmissibility *in vitro* (Zhang et al., 2020). The 25563-G-T (ORF3a:p.57Q>H) variant that has become dominant in USA is worthy of *in vitro* validation.

**Figure 2.**
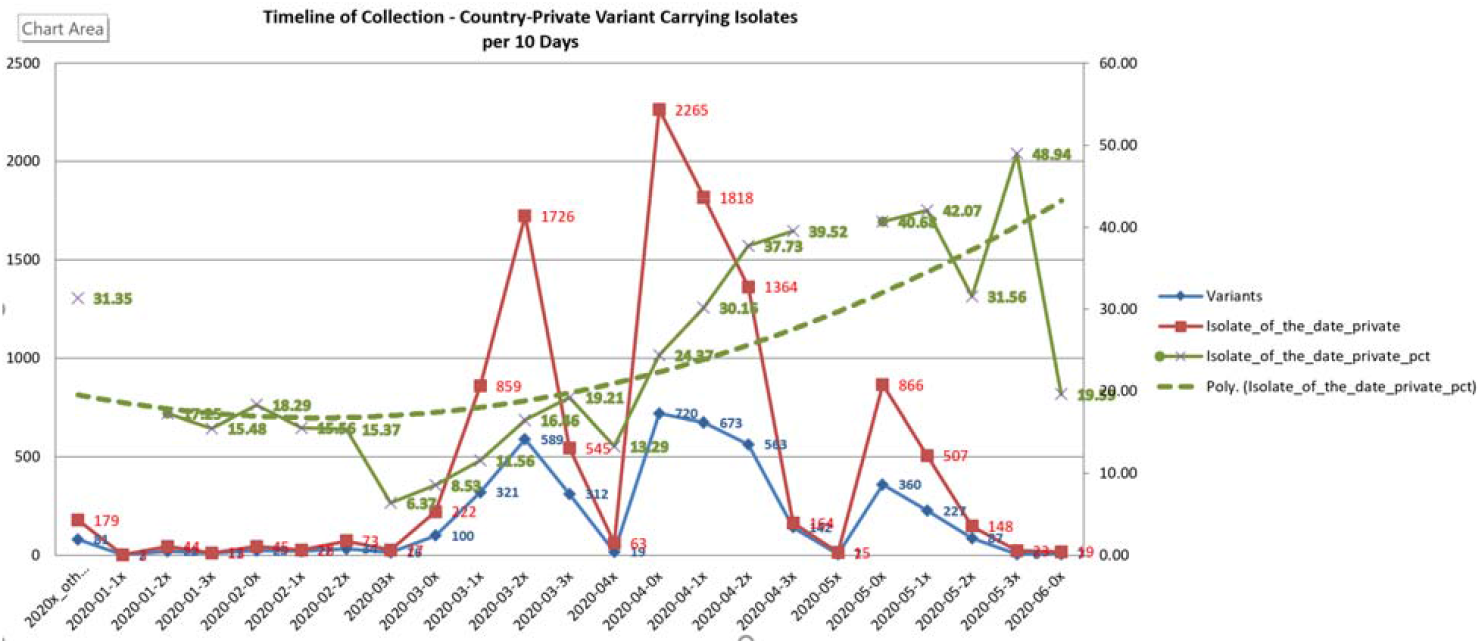
Timeline of collection - country-private variant carrying isolates. Date period with 10 or fewer total isolates were excluded. The trend line was polynomial at power of 2. Dates per 10 days were consolidated as a single time point.

**Figure 3.**
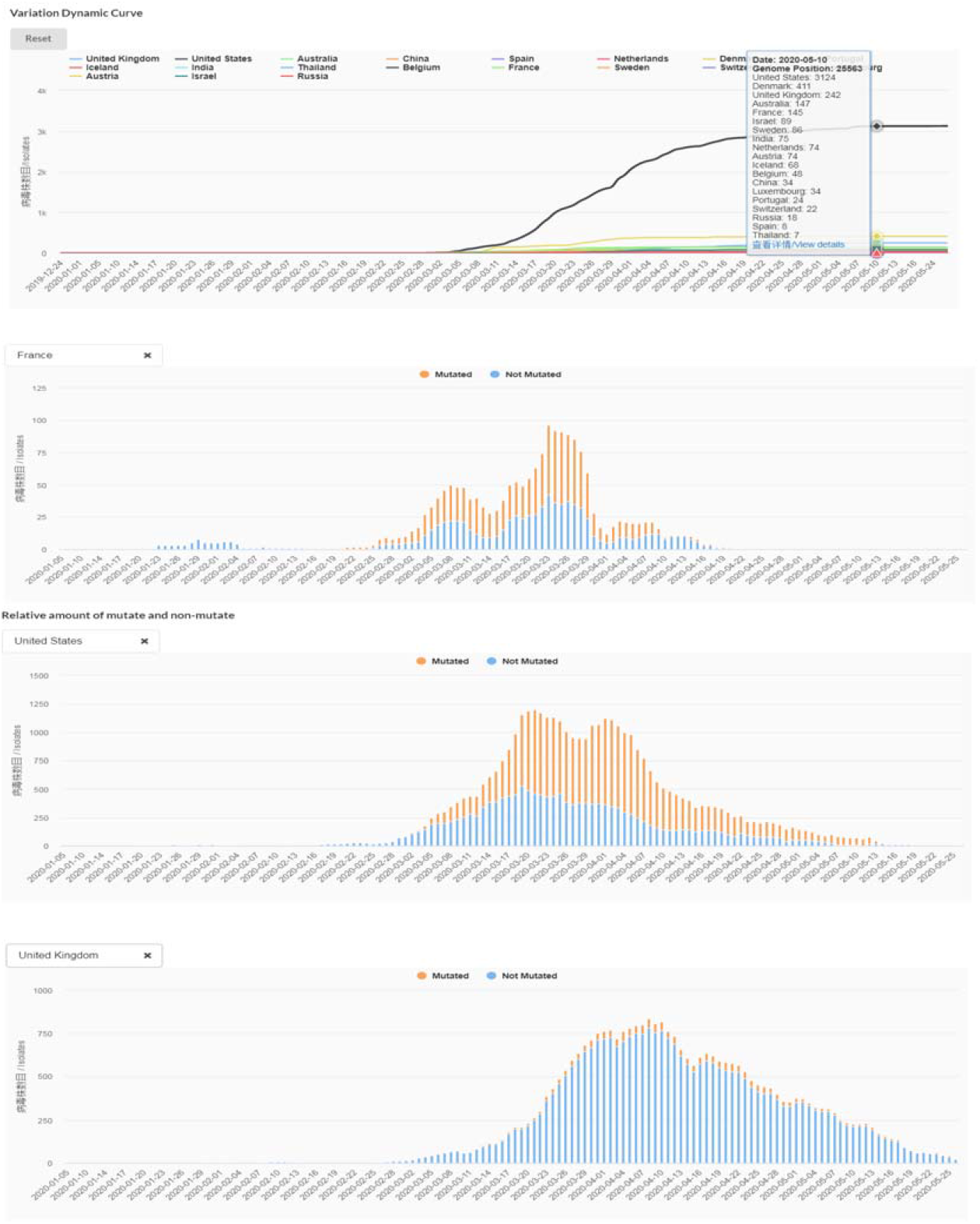
Variation dynamic curve and relative amount of mutate and non-mutate at position 25563. https://bigd.big.ac.cn/ncov/variation/annotation/variant/25563?lang=en

### Country-private variants

Twenty-six percent of the variants (1,583) present in at least 3 isolates were country-private variants (**Table 4**). Country-private variants, and similarly country-private haplotypes, were defined as variant where at least 98% of the variant-carrying isolates are from the same country and at most 2 isolates come from other countries no matter the total isolates. A single isolate could carry 2 or more country-private variants. They were identified in 10,238 isolates from 50 countries, with 25 countries having at least 10 isolates carrying the country-private variants. The UK had 896 private variants involving 6,142/22068 (28.8%) sequenced isolates whereas the USA had 329 private variants involving 2,166/10,402 (20.8%) sequenced isolates. Each of these private variants occurred in on average 30.0 isolates in the USA to 40.6 isolates in UK. Most other countries had less than 100 reported isolates carrying country-private variants. For instance, Australia had 52 private variants involving 428/2,169 (19.7%) of sequenced isolates and Iceland had 12 private variants involving 82 /596 (13.8%) sequenced isolates. This number is much lower than reported in a study comparing Iceland isolates against the 1,300-isolate GISAID March 22 collection (Gudbjartsson et al., 2020), because a larger 50k-isolate global collection was used for comparison in this study. Variants mentioned in that report may have strictly kept its Iceland-private status (i.e. 5142-C-T, gene orf1ab: p.Thr1626Ile; 25958-A-G, gene ORF3a:p.Tyr189Cys), or subsequently observed in other countries when more viral genome sequences were available (i.e. 24054-C-T, gene S:p.Ala831Val). Italy, a early COVID-19 epicenter has only one private variant 2269-A-T that was seen in 4 isolates.

By variant functional annotations, the majorities of country-private variants were missense variants totaling 896 variants involving 6,904 isolates, which were about 77% more frequent than the 586 synonymous variants affecting 3,893 isolates. High impact nonsense/stop gained variants affected only 86 isolates. For the genes affected, orf1ab ranked the first, as expected because of its large size, with 899 variants from 6,015 isolates, followed by S gene with 149 variants from 1,432 isolates. The N gene ranked the 3^rd^ with 80 variants in 515 isolates.

Regarding the emergence timeline of country-private variants and associated isolates, the numbers were low before mid-March when sequences genome numbers were less than 1,000. The new emergence peaked between mid-March and mid-April followed by a plateau that may be attributed to fewer genome sequences released since April 19^th^ (**Figure 2**). Smoothing the percentage of private variants at each 10-day interval revealed clear trend: 16.6% for January-February (206/1,244), 14.2% for March (3,379/24,777), 28.4% in April (5510/19,412), and the highest of 39.8% (1,578/3,967) in May to early June (**Figure 2**).

### Country-private haplotypes

Country-private haplotypes were identified in 39 countries when we limited it to haplotypes shared by at least 5 isolates, and required that at least 99% of the isolates were clearly from the same country and at most 2 isolates from other countries. A total of 802 haplotypes from 8,372 isolates were classified as country-private. Private haplotypes represented 22.4% (4,492), 16.6% (1,728) and 16.4% (356) of UK, USA, and Australia isolates, respectively.

Iceland and Thailand private haplotypes also represented 12.6% (75) and 33.7% (68) of the nation’s SARS-CoV-2 genome sequences, respectively. Iceland is geographically isolated from other countries. All members of the 74-isolate Iceland-private 6-variant haplotype “241-C-T, 3037-C-T,10323-A-G, 14408-C-T, 20268-A-G, 23403-A-G”, and the 34-isolate private 5-variant haplotype, “5142-C-T, 11083-G-T, 14805-C-T, 17247-T-C, 26144-G-T”, and all their 27 descendent isolates, were only seen from Iceland although they were present at about the same period of mid- to late-March 2020. Their potential immediate ancestral haplotypes with 5 and 4 variants were inferred with CHLA CARD Genome Tracker (Shen et al., 2020), and they were observed widely from Asia to Europe and America which was mostly sampled from late March. This may imply the importation of these haplotypes or their ancestors to Iceland, and then followed by localized outbreak without re-exportation to other countries.

Austria had a 36-isolate private haplotype “241-C-T, 3037-C-T, 12832-G-A, 14408-C-T, 23403-A-G, 28881-G-A, 28882-G-A, 28883-G-C” and its 2 Austria -private descendent haplotypes in 9 isolates. The isolates were collected from February 26 to late March. This haplotype’s earliest isolate originated from Italy (EPI_ISL_419656). The immediate ancestral haplotype was present in multiple European countries including Austria itself since February 24.

UK isolates had 464 private haplotypes from 4,942 isolates, which was 22.4% of all UK isolates. The 4 largest UK–private haplotypes were “1440-G-A, 2891-G-A, 28851-G-T” (UK-H1, 97 isolates) and its immediate descendant “1440-G-A, 2891-G-A, 28851-G-T and 1440-G-A, 2891-G-A, 25669-C-T, 28851-G-T” (UK-H2, 110 isolates), and “241-C-T, 3037-C-T, 10798-C-A, 13862-C-T, 14408-C-T, 23403-A-G, 28836-C-T” (UK-H3, 109 isolates), and “241-C-T, 3037-C-T, 14408-C-T, 22879-C-A, 23403-A-G” (UK-H4, 97 isolates). With CHLA CARD Genome Tracker and offline analysis, we found that the immediate ancestor of UK-H1 (2-variants, 1440-G-A, 2891-G-A) was widely circulating in multiple European countries including UK and Germany. All but one of the 986 isolates carrying these 4 haplotypes or any of their 403 immediate and remote descendent haplotypes were UK-private. Their evolutionary context was inferred from the closest ancestral and descendant isolates with Genome Tracker tool, and then combined with a randomly sampled representative global isolate collection for joint phylogenetic analysis. The result showed that these country-private clusters formed distinct clades (**Figure 4**). As the more comprehensive global summary, 700 of the 807 country-private haplotypes had 8,151 smaller decedent same-country private haplotypes each with fewer than 5 isolates per haplotype. The total isolates from them was 14,061, together with the major private haplotypes, 22,171 (45.8%) isolates carried country-private haplotypes globally. The percentage of isolates carrying private haplotypes were 29.9%, 28.2%, 46.4% and 59.6% in January-February, March, April and May 2020 respectively. It stayed low till March, but increased rapidly in April and May.

**Figure 4.**
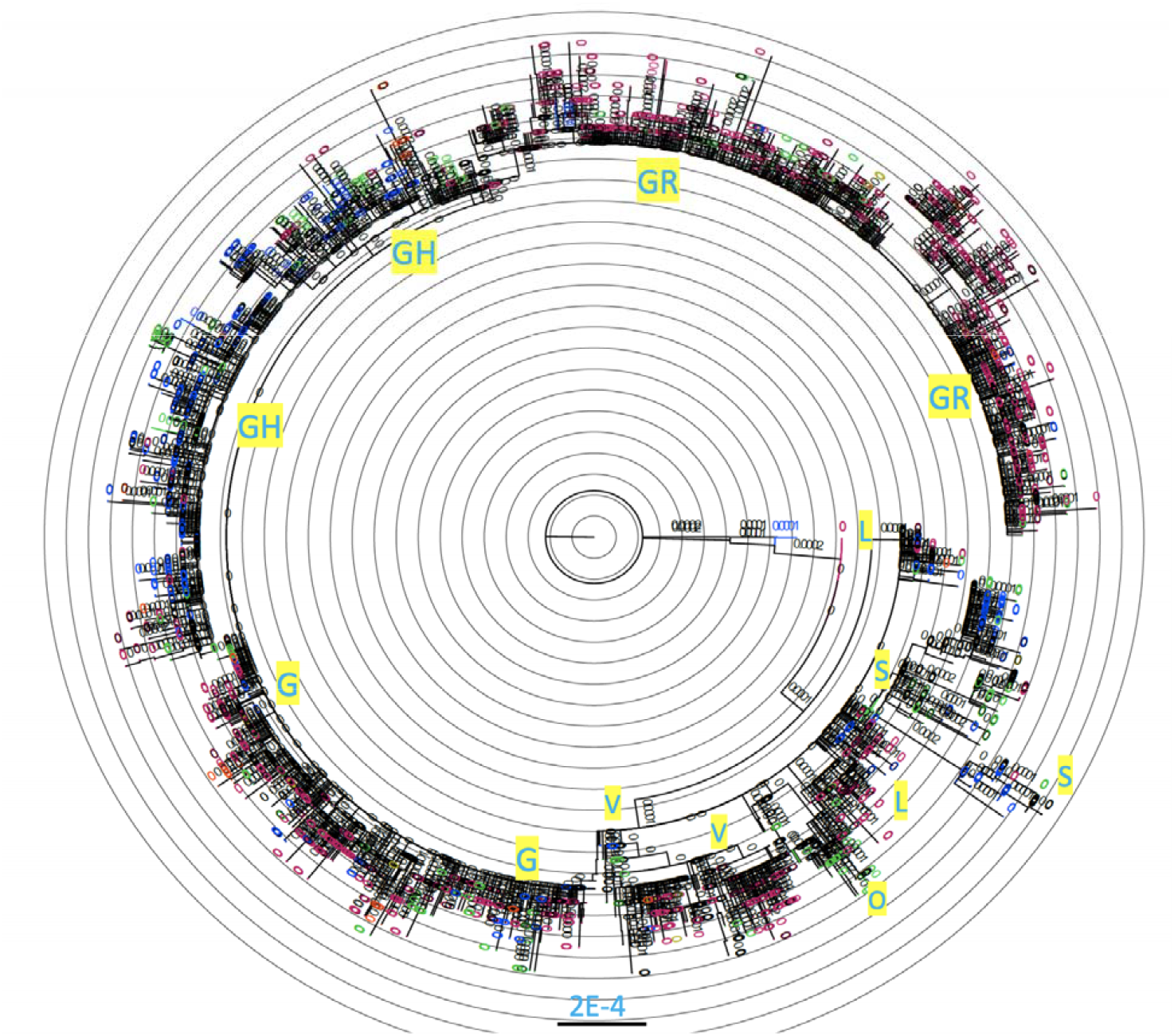
Maximum likelihood phylogenetic tree of representative isolates carrying Country-private or non-private haplotypes from the global isolates. Blue in yellow: Clade names. Red: UK-Private haplotypes, Blue: USA-Private haplotypes, Green: isolates with other Country-private haplotypes; Black: Non-country private haplotype isolates. 2E-4: distance scale bar; Rooted at the outgroup (MN996532|EPI_ISL_402131 bat/Yunnan/RaTG13/2013) and reference NC_045512 (MN908947|EPI_ISL_402125). Up to 6 isolates from each haplotype were included in the phylogenetic analysis. Each branch in the phylogenetic tree may represent a group of isolates.

There were relatively fewer private haplotypes or related isolates from Asia. It should be noted that the Asia private haplotypes were usually on the ancestral side, consisted of mostly 1 to 3 variants. This could reflect the earlier emergence and the gradual extinction of such private haplotypes as SARS-CoV-2 rates significantly declined in Asia. 21707-C-T was only seen in 21 isolates with China exposure and 1 Singapore isolate, and its identical and immediate descendant haplotypes were all sampled during the earlier phase (January to early February 2020) except for one China isolate EPI_ISL_454976 on March 4^th^, whereas most of the closest European or North American isolates carrying the 21707-C-T variant had 2 to over 10 extra variants which were evolutionarily remote and were sampled later between mid-March to late April, at least one month later. Similarly limited to a single country, the 8782-C-T haplotype was present only in 13 Singaporean isolates between Jan 27 to Match 2, but in no other country despite the fact that 3,965 other isolates across the world also carried this variant along with other variants.

### Geographic and temporal distributions of country-private haplotypes

Clear patterns of geographic specific distributions were evident at both variant and the haplotype levels from our stratified phylogenetic analysis. Geographic specific haplotypes were present in 8,372 of 20,414 isolates carrying haplotypes seen at least 5 times (41.0%). The UK, USA, Australia and Iceland together contributed 7,213 (86%) of the 8,372 isolates carrying country-private haplotypes. These countries are all geographically isolated from mainland Europe and Asia, where the initial outbreaks occurred. There were 19 private haplotypes that each had from 50 to 311 isolates. They were often continuously sampled across months but yet not “exported” to other countries despite the large numbers of isolates in circulation (over 20%). In contrast, only three country-private haplotypes with over 30 isolates were identified from the mainland European countries. One 35-isolate haplotype was from Austria, (“241-C-T, 3037-C-T, 12832-G-A, 14408-C-T, 23403-A-G, 28881-G-A, 28882-G-A, 28883-G-C”), the other one is a very young 36-isolate haplotype from Netherlands, only sampled between May 18 to 22 (“241-C-T, 1594-C-T, 3037-C-T, 11109-C-T, 13458-C-T, 14408-C-T, 22206-A-G, 23403-A-G, 25563-G-T, 27654-C-T”), and the third is Spain-private with 43 isolates (“8782-C-T, 9477-T-A, 14805-C-T, 25979-G-T, 28144-T-C, 28657-C-T, 28863-C-T, 29870-C-A”) and its 4 Spain-private descendent haplotypes. Other Country-private haplotypes from mainland European countries usually had lower numbers of isolates under 30 (**Table 6**). The UK-private haplotypes (Figure 4, colored in red) were mostly on different major clades than the USA-private haplotypes (Figure 4, colored in blue). This suggests reduced country-to-country transmissions potentially associated with travel restrictions.

**Table 5.** Functional annotation of country-private variants present in 3 or more isolates.

| **Annotation** | **Variants** | **Isolates** | **Isolates per private variant** | **Isolates purity** |
| --- | --- | --- | --- | --- |
| missense | 896 | 6904 | 43.2059 | 0.999629 |
| synonymous | 586 | 3893 | 17.3573 | 1 |
| intergenic | 74 | 485 | 28.303 | 1 |
| stop_gained; HIGH | 12 | 86 | 11.093 | 1 |
| frameshift; HIGH | 2 | 10 | 5.2 | 1 |
| downstream_gene,DISTANCE=55 | 2 | 7 | 3.5714 | 1 |
| downstream_gene,DISTANCE=63 | 1 | 3 | 3 | 1 |
| downstream_gene,DISTANCE=71 | 1 | 4 | 4 | 1 |
| downstream_gene,DISTANCE=86 | 1 | 6 | 6 | 1 |
| stop_gained&disruptive_inframe_insertion; HIGH | 1 | 3 | 3 | 1 |
| synonymous; inframe_deletion | 1 | 3 | 3 | 1 |
| downstream_gene,DISTANCE=88 | 1 | 3 | 3 | 1 |
| downstream_gene,DISTANCE=56 | 1 | 7 | 7 | 1 |
| downstream_gene,DISTANCE=32 | 1 | 3 | 3 | 1 |
| stop_lost&splice_var; HIGH | 1 | 13 | 13 | 1 |
| conservative_inframe_deletion; Moderate | 1 | 1 | 6 | 1 |
| inframe_deletion | 1 | 3 | 3 | 1 |
| coding_sequence | 1 | 4 | 4 | 1 |

**Table 6.**
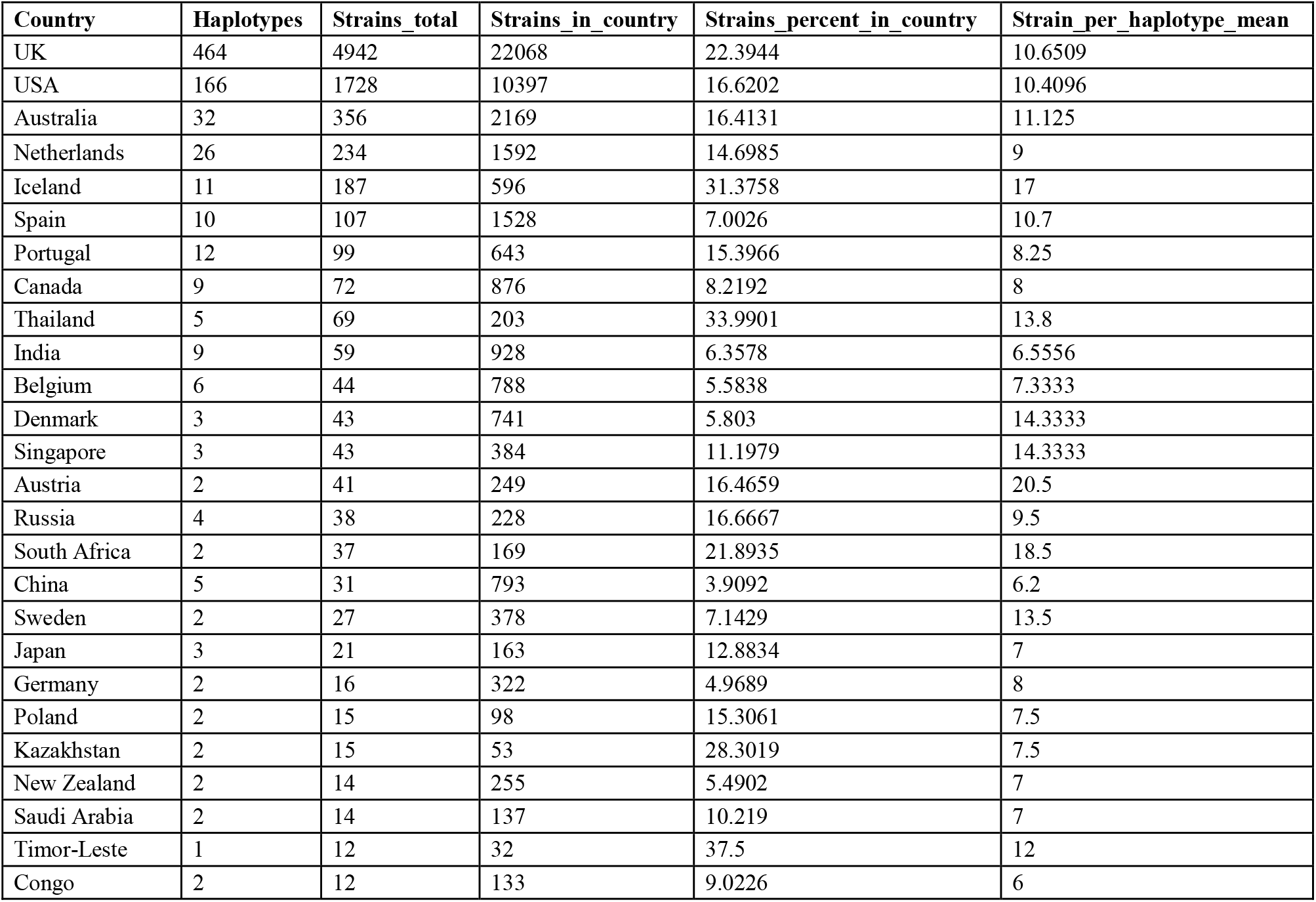
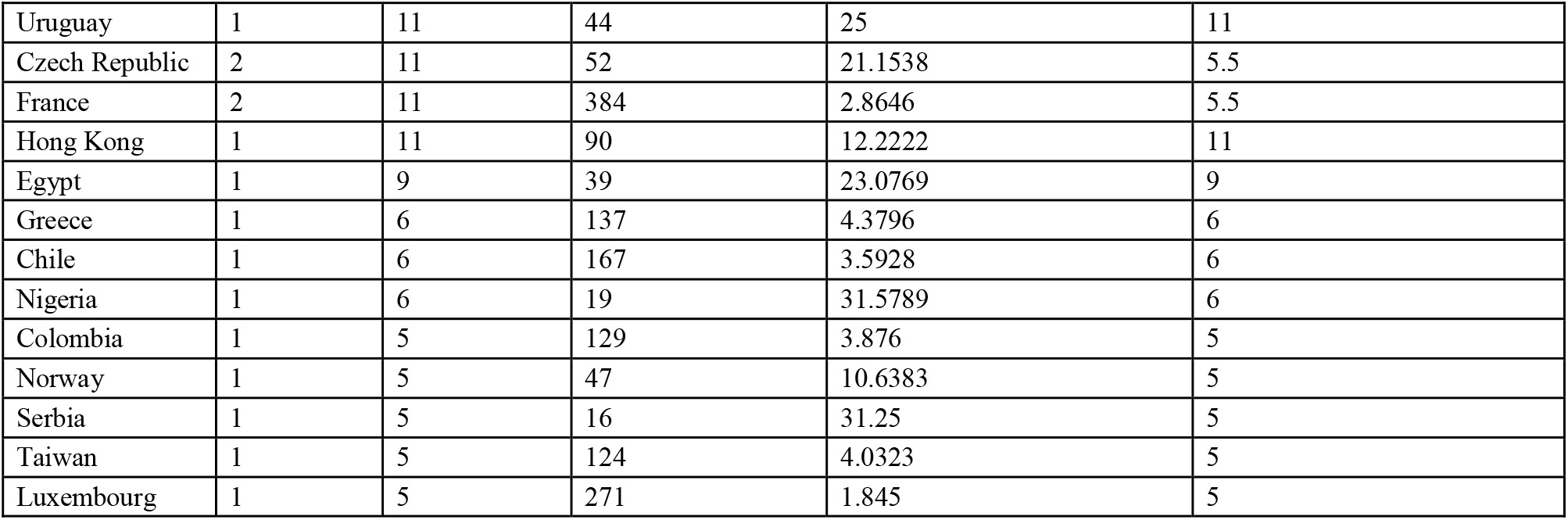
Country private haplotypes present in 5 or more isolates.

| Country | Haplotypes | Strains_total | Strains_in_country | Strains_percent_in_country | Strain_per_haplotype_mean |
| --- | --- | --- | --- | --- | --- |
| UK | 464 | 4942 | 22068 | 22.3944 | 10.6509 |
| USA | 166 | 1728 | 10397 | 16.6202 | 10.4096 |
| Australia | 32 | 356 | 2169 | 16.4131 | 11.125 |
| Netherlands | 26 | 234 | 1592 | 14.6985 | 9 |
| Iceland | 11 | 187 | 596 | 31.3758 | 17 |
| Spain | 10 | 107 | 1528 | 7.0026 | 10.7 |
| Portugal | 12 | 99 | 643 | 15.3966 | 8.25 |
| Canada | 9 | 72 | 876 | 8.2192 | 8 |
| Thailand | 5 | 69 | 203 | 33.9901 | 13.8 |
| India | 9 | 59 | 928 | 6.3578 | 6.5556 |
| Belgium | 6 | 44 | 788 | 5.5838 | 7.3333 |
| Denmark | 3 | 43 | 741 | 5.803 | 14.3333 |
| Singapore | 3 | 43 | 384 | 11.1979 | 14.3333 |
| Austria | 2 | 41 | 249 | 16.4659 | 20.5 |
| Russia | 4 | 38 | 228 | 16.6667 | 9.5 |
| South Africa | 2 | 37 | 169 | 21.8935 | 18.5 |
| China | 5 | 31 | 793 | 3.9092 | 6.2 |
| Sweden | 2 | 27 | 378 | 7.1429 | 13.5 |
| Japan | 3 | 21 | 163 | 12.8834 | 7 |
| Germany | 2 | 16 | 322 | 4.9689 | 8 |
| Poland | 2 | 15 | 98 | 15.3061 | 7.5 |
| Kazakhstan | 2 | 15 | 53 | 28.3019 | 7.5 |
| New Zealand | 2 | 14 | 255 | 5.4902 | 7 |
| Saudi Arabia | 2 | 14 | 137 | 10.219 | 7 |
| Timor-Leste | 1 | 12 | 32 | 37.5 | 12 |
| Congo | 2 | 12 | 133 | 9.0226 | 6 |
| Uruguay | 1 | 11 | 44 | 25 | 11 |
| Czech Republic | 2 | 11 | 52 | 21.1538 | 5.5 |
| France | 2 | 11 | 384 | 2.8646 | 5.5 |
| Hong Kong | 1 | 11 | 90 | 12.2222 | 11 |
| Egypt | 1 | 9 | 39 | 23.0769 | 9 |
| Greece | 1 | 6 | 137 | 4.3796 | 6 |
| Chile | 1 | 6 | 167 | 3.5928 | 6 |
| Nigeria | 1 | 6 | 19 | 31.5789 | 6 |
| Colombia | 1 | 5 | 129 | 3.876 | 5 |
| Norway | 1 | 5 | 47 | 10.6383 | 5 |
| Serbia | 1 | 5 | 16 | 31.25 | 5 |
| Taiwan | 1 | 5 | 124 | 4.0323 | 5 |
| Luxembourg | 1 | 5 | 271 | 1.845 | 5 |

### Clade and lineage assignment

The global isolates were assigned to major GISAID clades according to the diagnostic variant signatures and existing GISAID clades designations (**Table 7**). The largest clades were GR, G and GH clades. The smallest was the O clade. Examining the variants from the O clade isolates confirmed that there was no dominant variant or haplotype, where the most frequent haplotype (“6312-C-A, 11083-G-T, 13730-C-T, 23929-C-T, 28311-C-T”) presented in 3.6% isolates (77/2138), and the more frequent variants presented in the O clade isolates were 11083-G-T at 46.7%, 28311-C-T at 22.7%, and 13730_C-T at 20.4%. Isolates carrying the country-private haplotypes were less frequent for the O-clade (34.7%), but not far from other clades, between 40-50% among the haplotypes seen in 3 or more isolates. For per country clade ratios, USA has more GH (55.3%) and S (17.2%) clade isolates, whereas UK has more GR (44.2%) and G (25.3%) clades isolates but only 6.2% of GH and 1.1% of S clades (**Table 8**).

**Table 7.** The global and country private haplotypes and the clade present in >=3 isolates.

| **Clade** | **Signature** | **Haplotype / $\geq 3x$ /Global** | **Isolates / $\geq 3x$ /Global** | **Haplotype / $\geq 3x$ /Country private** | **Isolates / $\geq 3x$ /Country private** |
| --- | --- | --- | --- | --- | --- |
| G | 241-C-T, 3037-C-T, 23403-A-G | 535 | 5843 | 438 | 2772 |
| GH | 241-C-T,3037-C-T, 23403-A-G, 25563-G-T | 473 | 4999 | 381 | 2091 |
| GR | 241-C-T,3037-C-T, 23403-A-G, 28882-G-A | 722 | 7276 | 612 | 3878 |
| L | reference | 178 | 1345 | 143 | 854 |
| O | Other | 200 | 857 | 123 | 381 |
| S | 8782-C-T,28144-T-C | 175 | 1683 | 147 | 805 |
| V | 11083-G-T,26144-G-T | 184 | 1589 | 151 | 790 |

**Table 8.** The clade statistics for the 5 countries with over 1,000 isolate sequences.

| **Clade** | **Country** | **Isolates\*\*** | **%Isolates in country** |
| --- | --- | --- | --- |
| G | Australia | 556 | 25.63 |
| G | Netherlands | 550 | 34.55 |
| G | Spain | 639 | 41.82 |
| G | UK | 5578 | 25.28 |
| G | USA | 723 | 6.95 |
| GH | Australia | 452 | 20.84 |
| GH | Netherlands | 418 | 26.26 |
| GH | Spain | 40 | 2.62 |
| GH | UK | 1382 | 6.26 |
| GH | USA | 5751 | 55.31 |
| GR | Australia | 211 | 9.73 |
| GR | Netherlands | 299 | 18.78 |
| GR | Spain | 168 | 10.99 |
| GR | UK | 9746 | 44.16 |
| GR | USA | 393 | 3.78 |
| L | Australia | 32 | 1.48 |
| L | Netherlands | 208 | 13.07 |
| L | Spain | 10 | 0.65 |
| L | UK | 1787 | 8.10 |
| L | USA | 367 | 3.53 |
| O | Australia | 381 | 17.57 |
| O | Netherlands | 22 | 1.38 |
| O | Spain | 16 | 1.05 |
| O | UK | 550 | 2.49 |
| O | USA | 332 | 3.19 |
| S | Australia | 220 | 10.14 |
| S | Netherlands | 19 | 1.19 |
| S | Spain | 578 | 37.83 |
| S | UK | 252 | 1.14 |
| S | USA | 1793 | 17.25 |
| V | Australia | 105 | 4.84 |
| V | Netherlands | 73 | 4.59 |
| V | Spain | 57 | 3.73 |
| V | UK | 2773 | 12.57 |
| V | USA | 139 | 1.34 |

**\*\*:** Limited to the isolates from GISAID.

## Discussion

The generous and collaborative joint efforts by laboratories world-wide in generating and sharing SARS-CoV-2 genomes have made available a large amount of viral genome sequences and associated meta-data, for research, diagnosis, treatment development. We characterized the variants and haplotypes from all ~50,500 viral genome sequences available from multiple resources as of June 18, 2020. 6,070 variants were found to be present on 3 or more isolates, and 1,386 (26.8%) of them were country-private. The per gene variant density was quite consistent despite few hotspots. The comprehensive set of variants and haplotypes was integrated with demographic data, timeline of collection, and functional annotation, and made accessible at CHLA CARD web resource (Shen et al., 2020) for searching, browsing, and phylogenetic analysis. The global isolates were assigned to major GISAID clades (**Table 7**), but the clades were large with over 10,000 isolates in the G/GR/GH clades thus lacked resolution. There was an attempt to provide granularity for fine-scale lineage assignment (Rambaut et al., 2020), but a lineage classification system may need to handle the hierarchical structure with natural variant and haplotype profiles along the virus evolution as well as the reversal mutation and independent co-evolution. In our opinion, a system similar to the mitochondrial haplogroup such as modeled in the PhyloTree (http://www.phylotree.org) is worthy of consideration.

The travel restrictions between countries were mostly enforced in late March. March had the lowest percentage of isolates carrying country-private variants (14.2%) when compared to January and February (16.6%), and the percentage quickly doubled in April (28.4%) and tripled in May (39.8%) (**Figure 2**). SARS-CoV-2 may appears to have been spreading freely across countries through passenger traveling until late March, allowing for the spread of virus variant profiles to multiple countries. With the imposition of strict global travel bans in late March, the newly occurred variants were largely restricted to individual countries or even local regions as we observed in the US (Shen et al., 2020). It is concerning that over 40% of the recently sampled isolates carried country- or even province/state-private variants, which suggests a significant number of new variants are emerging and accumulating in a matter of months. The real-time monitoring of new viral genomes and *in silica* modeling of mutation effects are critical to timely identification of mutation similar to 23403A>G (S:p:614D>G) which quickly dominates and cause 4-time higher transmissibility *in vitro* (Daniloski et al., 2020; Zhang et al., 2020). It will be worthwhile to integrate the SARS-CoV-2 genotype and haplotype data with clinical data, and the national level transmission and morbidity data to search for potential highly transmissible haplotypes/genotypes or, conversely, high-morbidity but less-infectious ones. The host-virus genotype interaction possibility can also be assessed through multifaceted data integration. Such information can aid in the development of COVID-19 clinical management guided by both viral and host genotypes.

In summary, we performed an up-to-date and comprehensive variant and haplotype characterization of all reported SARS-CoV-2 sequences and isolates using the CHLA CARD (https://covid19.cpmbiodev.net/). Sign of positive and purifying selections were detected for further computational and experimental verification. There were a significant number of strongly localized outbreaks as evident from the country-private haplotypes and genotypes that involved multiple continents. Their locations, temporal transmissions, and timings of countrywide lock-down together demonstrated the effectiveness of existing travel restriction policies which may have been greatly enhanced by the natural geo-isolation, and the public health measures in controlling the transmission of SARS-CoV-2. The discovery of such a large number of country-private variants and haplotypes, in a matter of 6 months, raises legitimate concerns about the potential emergence of novel strains with increased virulence and/or transmissibility.

## Materials and Methods

### Source of SRAS-CoV-2 Sequences

The major external resources of SARS-CoV-2 isolates, genome sequences, and variants were GISAID, GenBank, CNCB, and NextStrain. Data was downloaded in various formats depending on the exact implementations of these external resources. Eighty-three viral sequences from the Center for Personalized Medicine, Children’s Hospital Los Angeles (CHLA) were also included. The viral genome sequences and metadata were cross-examined to identify and to remove duplicate entries present in multiple resources.

### Variant & Haplotype Analysis

Variants were called against the SARS-CoV-2 reference sequence (NC_045512.2) using dnadiff from MUMmer (Marçais et al., 2018, v.4). The analysis was automated with custom batch scripts. Variant functional annotation was carried out by snpEff (v4.31t). The ribosome slippage region positions in cDNA and protein were manually adjusted to account for the coronavirus programmed translational frameshift. The combination or constellation of all variants called in an isolate constitutes the haplotype of the isolate.

The isolates with excessive numbers of variants (>=30 per isolate) were excluded from the down-stream analysis as they likely represented low-quality sequences. All raw sequence, metadata, and analysis results were loaded into a MySQL database for integration and summarization.

### Phylogenetic Analysis

Evolution inference or relatedness analysis of viral isolates was carried out with full-length SARS-CoV-2 sequences using the Virus Genome Tracker tool of CHLA CARD (Shen et al, 2020). An isolate’s variant profile was compared against the global collection of over 50,500 viral sequences in CARD. The closest isolates with identical or the most similar variant profiles were identified and classified into identical, ancestral, or descendent categories.

For comprehensive and representative phylogenetic analysis to profile the over 50,000 viral genomes, 2 to 6 best quality sequences per haplotype were picked. More sequences were picked from the larger haplotypes. A total of 4,044 sequences were included in the final analysis. Leading and trailing missing bases “NNNN” were trimmed off genome sequences. Multiple sequence alignment (MSA) was done with MAFFT version 7.460 (Katoh et al, 2002; Katoh and Toh, 2008) using speed-oriented method FFT-NS-i (iterative refinement method, two cycles) optimized for large data-sets of thousands of full-length viral genomes. The original multiple sequence alignment (MSA) was manually examined in BioEdit to remove obviously low quality or outlier sequences. The resulting MSA was trimmed off 55 bases from the start, then 65 bases from the end to exclude these likely low coverage regions. Internal gaps over 5 bases in length or ones caused by missing bases were removed. The final trimmed MSA was used in MEGA-X (Kumar et al., 2018) for evolutionary inference via the Maximum Likelihood method assuming protein-coding sequences and under the General Time Reversible model (Nei and Kumar, 2000). Positions with over 2% alignment gaps, missing data, and ambiguous bases were excluded. The closest coronavirus sequence from bat (EPI_ISL_402131|MN996532, bat/Yunnan/RaTG13/2013) was included in MSA and tree building as the outgroup to root the tree. The phylogenetic tree was visualized in MEGA X and FigTree v1.4.4 (http://tree.bio.ed.ac.uk/software/figtree/).

### Categorical Analysis

Variants and haplotype were labelled and stratified with isolate metadata, including residential and exposure counties, along with the date of sample collection, sequence technology and quality, and other demographic data such as patient gender and age.

Enrichment tests among groups defined by meta-data were carried out with the Chi-square test. For missense vs synonymous variant analysis, the in-silica predicted missense and synonymous variant numbers were used as the background assuming neutral selection on all coding sites. Then the observed variant and isolated numbers were used in the Pearson’s Chi-squared test with Yates’ continuity correction to identify enrichment and depletion by function types.

The clade assignments from the GISAID.org were included when available, and used in stratification of the haplotypes from the matching isolates (**Table 7**). The clades were G, GH, GR, L, O, S and V clades.

## Acknowledgements

We are thankful to other staff members of the CHLA Virology Laboratory and the Center for Personalized Medicine laboratory for their strongest support of this project, while providing patient care and battling the COVID-19 pandemic together. The SARS-CoV-2 genomes and meta data were generously shared via GISAID, GenBank, China National Center for Bioinformation (CNCB), Nextstrain and other sources. We gratefully acknowledge the originating and submitting laboratories for making the SARS-COV-2 sequences and associated metadata available via these public resources. For a full list of such contributors, please see from “Virus Isolate” tool on CHLA CARD (https://covid19.cpmbiodev.net/) the full lists of “Originating labs”, “Submitting labs” and “Authors”.

## Notes

https://covid19.cpmbiodev.net/

### Competing Interest Statement

The authors have declared no competing interest.

